# *Flux Transformer* learns intracellular metabolic fluxes from isotope labeling and measured reaction rates

**DOI:** 10.64898/2026.11.16.767180

**Authors:** Daniel R. Weilandt, Md Toki Tahmid, Rachid Ounit, Jorg Schwender, Mona Singh, Joshua D. Rabinowitz

## Abstract

Cells convert incoming nutrients into energy and biomass via metabolic fluxes. Quantifying these fluxes involves feeding cells isotope-labeled nutrients, quantifying metabolite labeling patterns, and inverting the labeling data to retrieve the underlying flux distribution. Traditional metabolic flux analysis (MFA) is both computationally expensive and frequently requires manual refinement of the cost function to obtain plausible results. For this reason, MFA has remained inaccessible to non-experts. Here we present *Flux Transformer*, an AI model that uses reaction-specific attention to integrate isotope-labeling data with available reaction-rate measurements, reconstructing mass-balanced steady-state fluxes. The model is trained on simulated data with added noise and identifies the underlying fluxes from these data more effectively than traditional MFA. Provided experimental data from wild-type and mutant bacteria, yeast, plants, and mammalian T cells, it efficiently and accurately recovers expert-curated MFA fluxes. By making flux analysis accessible, *Flux Transformer* is poised to accelerate metabolic research and engineering.

## Main

Life relies on the continuous transformation of nutrients into energy and biomass through a network of interconnected biochemical reactions. Although the architecture of this network is largely encoded in the genome, the cellular phenotype is determined by the rates (fluxes) at which metabolites are processed by the biochemical reactions. These metabolic fluxes quantify how cells allocate resources, generate energy, and synthesize molecular building blocks.

Some metabolic fluxes can be measured directly. In cultured cells, for example, nutrient uptake and byproduct secretion rates can be quantified from the change in metabolite concentrations in the medium over time. But intracellular fluxes cannot be directly quantified. The most established way to determine them is through isotope-tracing experiments combined with metabolic flux analysis (MFA)^1,2^. In this approach, cells are supplied with isotopically labeled nutrients, and the resulting labeling patterns of intracellular metabolites are measured by nuclear magnetic resonance^3^ or mass spectroscopy^4^. The labeling patterns are then used to deduce the underlying fluxes.

Determination of fluxes from labeling patterns is a challenging inverse problem. Computationally, isotope labeling patterns can be directly computed from flux distributions using approaches including elementary metabolite units (EMU)^5^. In contrast, the inverse problem of determining flux distributions from labeling patterns does not have a direct analytical solution. Instead, it is solved by serially calculating the isotope labeling patterns generated from a candidate flux distribution and using non-linear optimization methods to converge on a flux distribution that best fits experimental data using a cost function.^6^ One challenge is that input data are heterogeneous, consisting of labeling patterns and measured rates (e.g., metabolite exchange rates with the media or rates of biomass formation). As a result, the fitting process is commonly not robust and instead sensitive to the exact weights put on different data, with expert modulation of the weights (or other aspects of the fitting process) required to return plausible fluxes. Moreover, the nonlinear optimization process is computationally expensive and does not guarantee convergence to the global optimum. Collectively these factors make MFA inaccessible for typical biologists.

Machine learning has transformed problems across biology, from protein structure prediction to single-cell analysis. Early efforts used machine learning to identify, from tracing data, relevant flux-related features but did not actually determine fluxes^7,8^. Further efforts trained surrogate models to approximate the forward problem and identified fluxes by fitting the predictions of the surrogate model to the experimental data^9,10^. Most recently, a two-stage machine learning framework combined imputation with prediction to infer relative fluxes from tracer data^11^. This method, however, did not address the most common real-world MFA context: Both isotope labeling data and selected reaction rates are experimentally measured and must be integrated to yield a unified accurate flux distribution.

Here present *Flux Transformer*, an AI model that uses an attention mechanism to directly map all relevant available data (isotope tracing data and selected reaction rates) to the underlying flux distribution. Simulated data generated from a diverse range of physiologically realistic flux distributions, with added noise, were used to train the AI model to infer the underlying flux distribution from the given measurements. Applying this strategy to data from five organisms (*Escherichia coli, Saccharomyces cerevisiae, Issatchenkia orientalis, Thlaspi arvense L*., and *Mus musculus*), we find that *Flux Transformer* more accurately recapitulates the full flux distributions underlying noisy simulated data than traditional MFA. Moreover, *Flux Transformer* reproduces expert-curated MFA flux estimates from experimental data without the manual refinement of the cost function or other aspects of the fitting process commonly required in MFA.

### *Flux Transformer:* Attention-based flux inference with steady-state constraints

*Flux transformer* frames flux identification as a machine-learning problem. For training data, we simulate isotope labeling patterns based on a large set of physiologically reasonable flux distributions. Rather than iteratively refining the flux distribution to match the predicted labeling as in traditional MFA, an AI model inverts the relationship between the simulated data and the flux distribution. It does so by extracting generalizable input data patterns that inform the underlying flux values (Fig. 1a).

**Figure 1.**
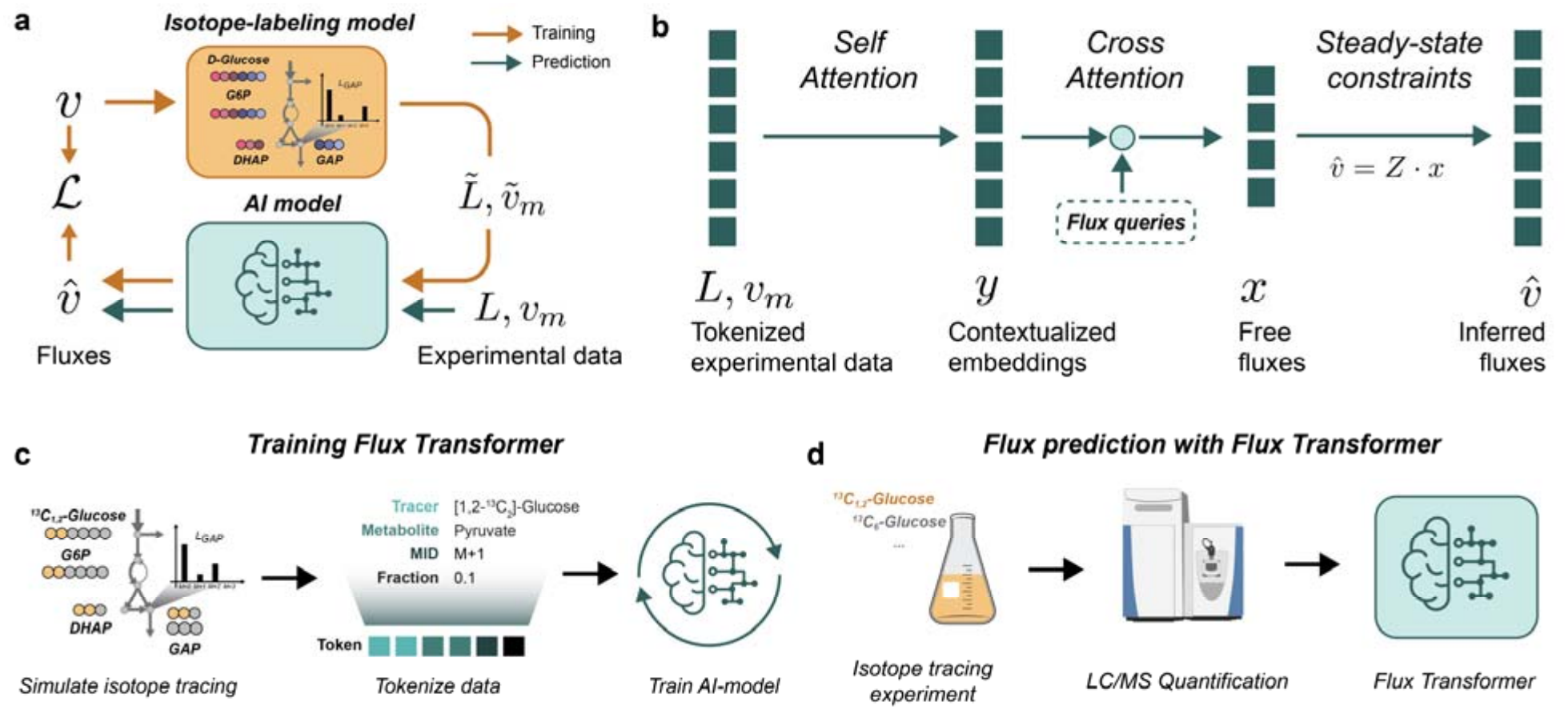
*Flux Transformer* infers cellular metabolic fluxes from isotope labeling data and experimentally measurable rates. (a) Overview of the approach. During training (orange), simulated data (labeling patterns 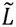 and experimentally measurable rates 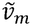) are generated from physiological fluxes 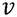, adding appropriate noise. An AI model then learns to invert these data, by minimizing the loss ℒ between and the output fluxes 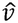 and the ground truth fluxes *v*. During prediction (teal), the trained model determines fluxes 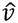 from experimental data (*L*, v_*m*_). (b) *Flux Transformer* architecture. Tokenized measurements are contextualized by the self-attention layers of a transformer encoder, yielding *contextualized embeddings y*. Learned *flux queries* cross-attend to these embeddings, producing a set of *free fluxes x*, which map to the full predicted flux distribution 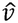, via the basis *Z* spanning the steady-state flux space. (c) Training workflow: (i) Isotope tracing experiments are simulated for diverse physiologically reasonable flux profiles and appropriate noise is added, (ii) Simulated isotopologue and rate measurements are tokenized, and (iii) Tokenized data is passed to train *Flux Transformer*, which is trained by minimizing the difference between flux predictions 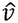 and “ground truth” *v*. (d) Flux identification from experimental data: (i) Cells are cultured with isotopically labeled tracers, (ii) labeling patterns and measurable rates are quantified, and (iii) after translating these measurements into tokens *Flux Transformer* identifies full flux distributions including uncertainty.

A challenge in machine learning is how to encode input data so that an AI model can efficiently extract the information relevant to its task. This is especially true when inputs combine different data types. Input data for isotope-based flux analysis consists of two data types: (i) labeling patterns and (ii) a small set of directly measured reaction rates. To encode these two data types, *Flux Transformer* represents each observation together with a compact description of its biological identity. Each isotope labeling measurement is represented as a typed token^12^ encoding its metabolite identity, mass shift, tracer identity, and measured labeling fraction. Experimentally measured fluxes are represented by a token encoding their reaction identity and flux value. This converts the heterogeneous measurements used in MFA into a common, structured representation (token), without discarding the biological meaning of each observation.

Because individual labeling observations are insufficient to determine fluxes, the information content of the tokens needs to be integrated. An effective way to do this is self-attention, which updates each token with a weighted average over all tokens in the input, giving more weight to those most relevant to the token being updated. What counts as relevant is determined by parameters learned during training^13^. Passing the tokens through a series of self-attention layers lets each measurement be interpreted in the context of the full experiment (Fig. 1b). As self-attention operates on inputs of arbitrary size, this architecture naturally handles missing data, which is a common occurrence in isotope tracing experiments.

The model must ultimately output a quantitative flux value for every reaction. To do so efficiently, we exploit the fact that biological systems typically operate at metabolic pseudo-steady state: Intracellular metabolite concentrations remain nearly constant, with the fluxes producing each metabolite roughly balancing with those consuming it. This imposes a set of linear mass-balance constraints on the fluxes. Based on this constraint, the full flux distribution can be expressed as a linear combination of a small set of free fluxes (Fig. 1b). Accordingly, the model only needs to predict a few free fluxes. *Flux Transformer* uses cross attention to compute the free fluxes from the contextualized embeddings. Cross-attention works like self-attention except that, instead of the input tokens, a set of learned queries is updated with weighted averages over the inputs (here contextualized embeddings), giving more weight to those most relevant to each query’s part of the prediction (Fig. 1b)^14^. Whereas self-attention gives each measurement meaning in the context of the others, cross-attention interprets the contextualized data to predict the fluxes.

Building a new *Flux Transformer* model from an existing metabolic network follows three steps (Fig. 1c): (i) generating synthetic labeling data for a large set of physiological flux distributions (with realistic noise added), (ii) tokenizing the data, and (iii) training the model. Training is based on minimizing the error between the learned and “ground truth” fluxes (i.e. the fluxes used to generate the synthetic labeling data). Rather than training a single model, we train an ensemble of models from different random initializations. Because each model converges towards a slightly different solution, their predictions agree where the data tightly constrain the fluxes and diverge where they do not, so the ensemble provides both a flux estimate and a measure of its uncertainty^15^.

Once trained for a given reaction network, the ensemble predicts fluxes directly from tokenized experimental data (Fig. 1d). We report the medoid of the ensemble predictions (flux distribution with the smallest total distance to all others) as the best flux distribution estimate. To quantify uncertainty, we repeatedly draw random sub-ensembles from the full ensemble and compute the medoid of each. Confidence intervals for the flux estimates are then taken from the percentiles of the resulting distribution that correspond to the desired confidence level. For example, a two-tailed 95% confidence interval (significance level of 5%) uses the 2.5^th^ and 97.5^th^ percentiles. In this manner, *Flux Transformer* outputs mass-balanced flux distributions with error estimates.

## Results

### Flux Transformer accurately identifies metabolic flux

We set out to test if *Flux Transformer* can recover intracellular fluxes from noisy data. To this end, we examined metabolic models of (i) *E. coli* ^16,17^, (ii) *S. cerevisiae*^18^, (iii) *I. orientalis*^19^, (iv) pennycress embryos^20^, and (v) mouse T cells^19^. To train the *Flux Transformer* models, we generated synthetic data that mirrored physiological fluxes of the relevant organisms and that was restricted to metabolites reliably detected and identified experimentally (Supplemental Data S2, Tables 1 and 2). We then used Monte Carlo simulation to generate 10,000 steady-state flux vectors for each organism and trained an ensemble of 20 AI models as described above. To test their performance, we withheld 1,000 of the generated flux vectors from training and added 5% relative noise to reaction rate measurements and 1% absolute measurement noise to each labeling measurement (see Methods).

We measured the performance of *Flux Transformer* by comparing inferred fluxes to the ground truth, computing R^2^ reaction by reaction across the 1,000 withheld flux vectors (Fig. 2a,b). Key modifications to the model formulation and to the training processing improved performance as compared to earlier simpler versions of the model, as we demonstrate via ablations. Two modifications relate to the distinction between the linearly independent free fluxes and the associated full flux distribution. In principle, the full flux distribution can be calculated from any set of linearly independent fluxes that span the mass-balanced (i.e. steady-state) flux space. In practice, however, the choice of free fluxes is crucial to model performance. Specifically, it is essential for any directly measured reaction rate to correspond to a free flux (Fig. 2c). Otherwise, the model struggles to balance the other relevant free fluxes to match the experimental rate measurement. Also, training is more effective when the objective is matching the small set of free fluxes rather than comparing output to the full flux distribution (Fig. 2d). We also found that penalizing the difference between data and prediction loss function by the variance in the dataset was crucial for performance (Fig. 2e; Methods). Most importantly we could show that basic neural network architecture (multi-layer layer perceptron; no tokenization) with a similar number of trainable parameters was not able to achieve the same performance as *Flux transformer* (Fig. 2f).

**Figure 2.**
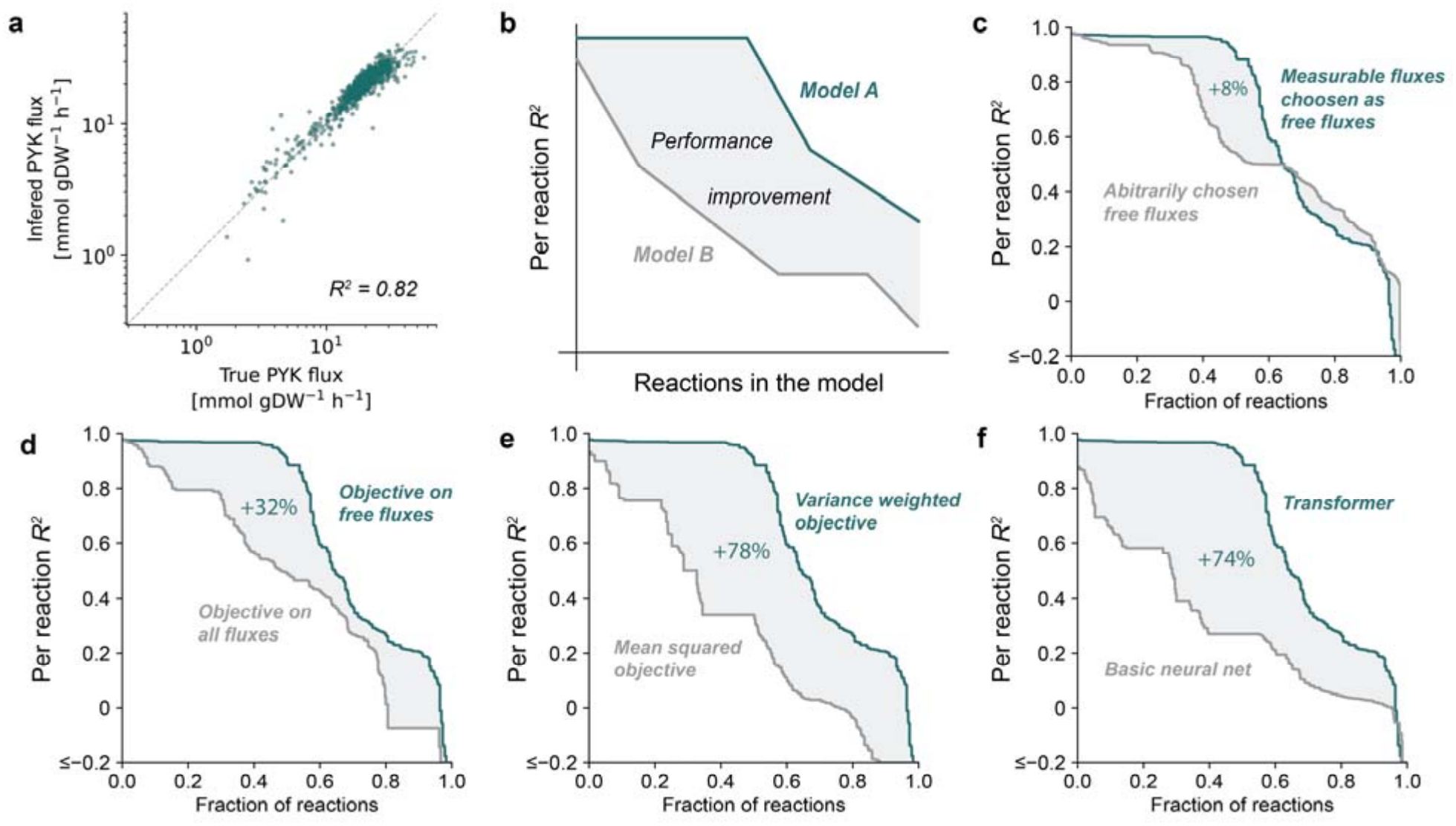
Systematic optimization of *Flux Transformer* improved prediction accuracy. (a) Comparison of input (“true”) to inferred fluxes for the pyruvate kinase reaction in *I. orientalis* (R^2^ = 0.82). Each point represents the *Flux Transformer* output for the PYK reaction given a complete input dataset, and the dashed line indicates the identity relationship. (b) Schematic illustrating how improvements in model design can be quantified at the reaction level. Reactions are ranked by predictive performance (R^2^), and the shaded region represents the aggregate performance gain of one model relative to another. (c–f) To identify design choices that maximize predictive accuracy, we compared alternative model configurations using the approach in (b). Percentage numbers indicate relative change in area under the curve. Selecting experimentally measurable fluxes as free fluxes (c), defining the training objective on free fluxes rather than all fluxes (d), using a variance-weighted objective (e), and adopting a transformer architecture (f) each improve prediction accuracy across the reaction network. Curves show ranked per-reaction R^2^ values, with reactions ordered from highest to lowest performance.

We next compared the overall performance of *Flux Transformer* to traditional MFA. Initially we planned to benchmark *Flux Transformer* against two MFA software packages (^13^C-FLUX v3.0 and INCA v2.5). We found that, for simulated data from the *I. orientalis* metabolic network, ^13^C-FLUX reached acceptable solutions with a single start (median R^2^ > 0.9), while INCA required at least 1000 restarts to reach a similar level of performance (median R^2^ ≈ 0.9; Fig. S1a). We therefore only provide comprehensive benchmarks against ^13^C-FLUX. We found that *Flux Transformer* resolved more reactions (R^2^ > 0.9) than traditional MFA (Fig. 3a-f), and returned more accurate fluxes overall (Fig S1b). *Flux transformer* also identified fluxes orders of magnitudes faster than ^13^C-FLUX and INCA (Fig. S1c). Moreover, machine learning approaches that rely solely on labeling patterns without incorporating measured fluxes (e.g. ML-Flux^11^) exhibited less reliable flux recovery (Fig. S1d).

**Figure 3.**
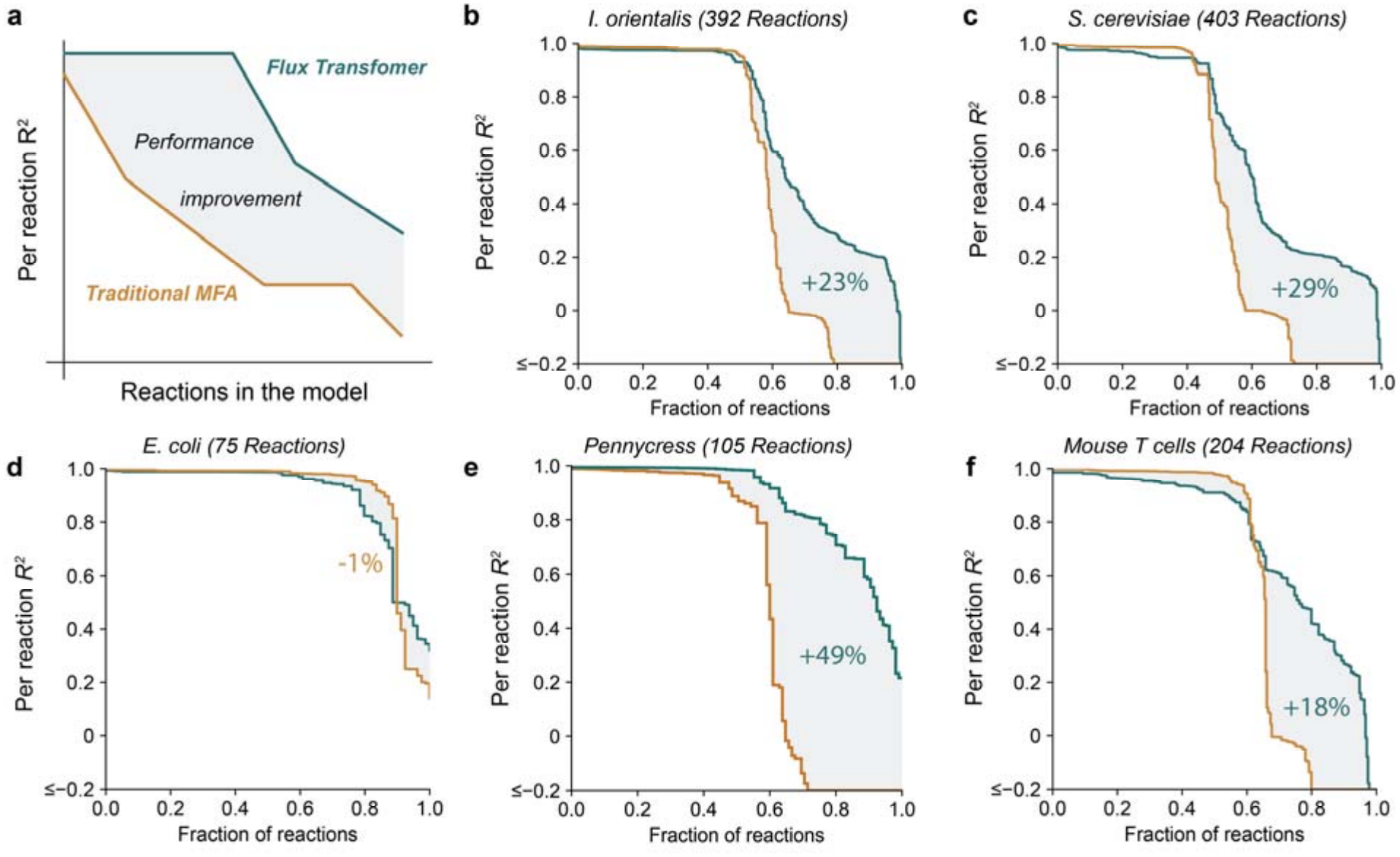
*Flux Transformer* efficiently resolves more fluxes than traditional MFA. (a) Schematic illustrating the comparison of reaction-level performance between *Flux Transformer* and traditional metabolic flux analysis (MFA). Reactions are ranked by accuracy of the inferred fluxes (R^2^). The shaded region represents the aggregate performance gain achieved by *Flux Transformer*. (b–f), To evaluate generalizability across biological systems and network sizes, we compared reaction-level flux inference for *Flux Transformer* and traditional MFA for the metabolic networks of *I. orientalis* (b), *S. cerevisiae* (c), *E. coli* (d), pennycress embryos (e), and mouse T cells (f). *Flux Transformer* consistently matched or increased predictive accuracy. Percentage numbers indicate relative change in area under the curve.

### Flux Transformer recovers expert curated fluxes form experimental data

We next tested whether *Flux Transformer* performs equally well on experimental data. To do so, we collected examples of published expert-curated MFA results and the underlying experimental data^19,21^. These experiments used uniformly and positionally labeled glucose, as well as uniformly labeled glutamine as tracers. Specifically, Park *et al*. traced into *E. coli*, with [U-^13^C_6_]glucose, [1,2-^13^C_2_]glucose, and [3-^13^C_1_]glucose, data from Shen *et. al* used [U-^13^C_6_]glucose, [1,2-^13^C_2_]glucose in both two different yeast species and in mouse T cells, and additionally [U-^13^C_5_]glutamine in the mouse T cells, whereas Shi et. al. used a mix of [U-^13^C_6_]glucose and [1,2-^13^C_2_]glucose, as well as [2,3-^13^C_2_]alanine and [1-^13^C_1_]alanine in pennycress embryos^20^. We provided *Flux Transformer* with the reported labeling and exchange rate measurements. It then inferred fluxes. We then used the fluxes which it determined to compute isotopolgoue fractions, and we evaluated flux and labeling predictions against the published data (Fig. S3a). Across all four organisms, *Flux Transformer* recovered both the measured labeling patterns (median R^2^ = 0.82; Fig. S3a) and the expert-curated MFA fluxes (median R^2^ = 0.90; Fig. 4b). This demonstrates that although flux transformer was trained exclusively on simulated data, it learned mappings between the data and the fluxes that transfer to real measurements.

**Figure 4.**
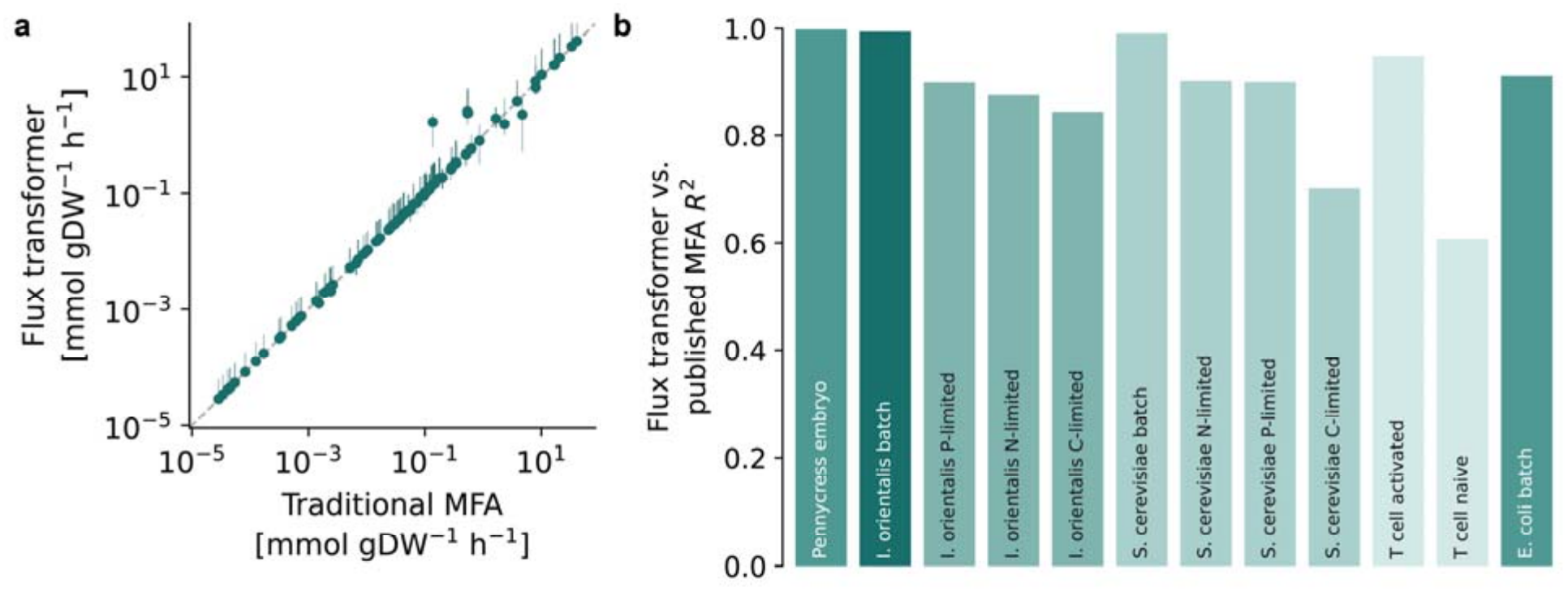
*Flux Transformer* recovers expert-curated fluxes from experimental data across diverse biological systems. (a) *Flux Transformer* inferred fluxes versus published, expert-curated metabolic flux analysis (MFA) flux estimates. Only reactions which MFA could reliably quantify in the validation dataset (R^2^ > 0.9) were compared. Each point represents a reaction flux; dashes mark the line of identity. (b) Agreement between *Flux Transformer* inferred fluxes and published MFA fluxes across benchmark datasets. Bars report R^2^ across all flux values within each dataset.

In batch cultures of *E. coli, S. cerevisiae*, and *I. orientalis*, fluxes quantified with *Flux Transformer* agreed over five orders of magnitude with those reported by experts (R^2^ > 0.9; Fig. 4a; Fig S2a-c). For *E. coli*, this held even though *Flux Transformer* was trained on a different network model^16^ than the one used by Park et al., indicating that *Flux Transformer* has learned biochemical logic rather than features of a specific network model. Importantly, the reactions on which *Flux Transformer* and expert MFA disagreed were those Flux Transformer reported with the widest confidence intervals (Fig. 4a; Fig. S2a-l), showing that the model has learned to recognize which fluxes it cannot determine reliably. Consistent with this, the datasets showing the greatest disagreement between expert MFA and Flux Transformer predictions, namely carbon-limited *S. cerevisiae* and naïve T cells, also exhibited poor agreement between predicted and measured labeling patterns for both *Flux Transformer* and expert MFA (Fig. S4). Thus, in contexts where fluxes are well defined by the data, *Flux Transformer* recapitulates expert-curated MFA fluxes, and in contexts where data fail to reliably define the fluxes, *Flux Transformer* properly returns wide confidence intervals.

### Flux Transformer accurately and efficiently predicts fluxes in mutant cells

Metabolic flux analysis is widely used in metabolic engineering to assess how genetic perturbations reroute metabolism. As a final test, we therefore asked whether *Flux Transformer*, trained only on simulated wild-type flux distributions, can predict fluxes in mutant cells. We sourced public ^13^C-labeling data quantified by GC-MS, for wild-type *E. coli* and 20 single-gene deletion strains of central carbon metabolism, each previously analyzed by MFA^22^ (Fig. 5a). We then retrained the *E. coli* model based isotopologues of GC-MS fragments instead of intact isotopologues of intact metabolites. As before, the model was given only the labeling and reported rate measurements but nevertheless recovered the published fluxes across all strains (R^2^ ∼ 0.95; Fig. 5b). It is noteworthy that, as with the comparison to the Park *et al*. data^21^, we here also used a different network model^16^ for training than Long and Antoniewicz did for their MFA^22^, yet recovered excellent agreement.

**Figure 5.**
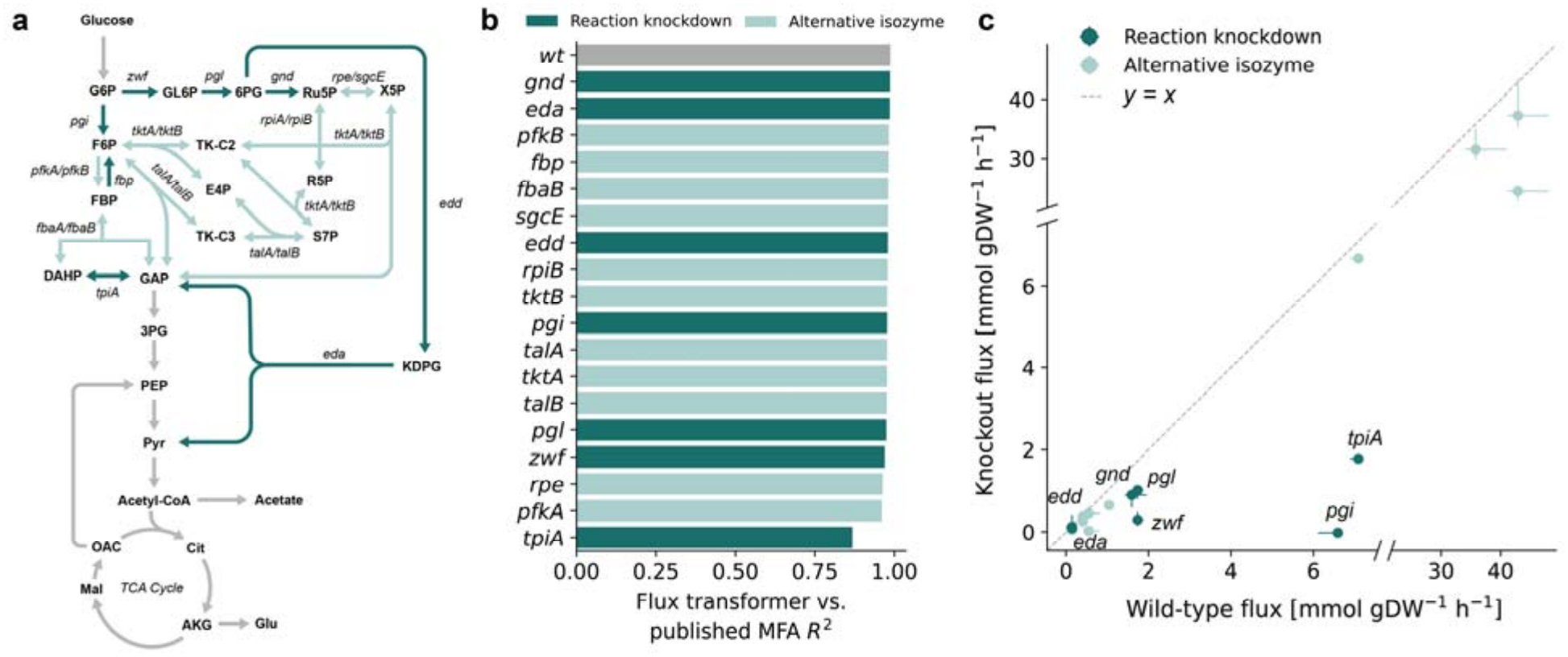
*Flux Transformer* correctly infers fluxes in *E. coli* knockout strains. a) Schematic map of central carbon metabolism of *E. coli* highlighting reactions impacted in knockout strains, dark green indicates reactions where the knockout removes the reaction, light green indicates cases where at least one alternative isozyme is present^22^. b) Agreement between *Flux Transformer* inferred fluxes and published metabolic flux analysis (MFA) fluxes for individual genetic perturbations. Bars report R^2^ across all flux values for each genetic perturbation. c) Fluxes in perturbed strains versus the corresponding wild-type fluxes. Dark teal points denote reactions where the only available enzyme is genetically perturbed and light teal points denote reactions with an alternative isozyme available. Error bars indicate uncertainty estimates from *Flux Transformer*, and the dashed line indicates equal flux in the wild-type and perturbed strains (y = x).

Knockout of certain genes, such as enzymes that are not directly involved in the measured network or one of multiple isozymes catalyzing the same reaction, change fluxes only modestly (Fig. 5b; light bars). Loss of core network enzymes, however, such as phosphoglucose isomerase (pgi), triosephosphate isomerase (tpiA), 6-phosphogluconate dehydrogenase (gnd), 6-phosphogluconolactonase (pgl), glucose-6-phosphate dehydrogenase (zwf), 6-phosphogluconate dehydratase (edd), and the aldolase of the Entner-Doudoroff pathway (eda) trigger massive metabolic flux rerouting (Fig. 5b; dark bars). In both cases, *Flux Transformer* captured knockout phenotypes accurately. Only the fluxes for the triosephosphate isomerase knockout agreed less well with MFA results (tpiA; R^2^ = 0.85). Notably, MFA using the standard *E. coli* network also performed poorly for this strain, and Long and Antoniewicz adopted an altered network model for this strain specifically^22^, a modification we did not adopt.

Examining the deleted reactions themselves revealed that *Flux Transformer* consistently predicts low fluxes for genetically ablated reactions, although it was never trained to perform this task (Fig. 5c). For phosphoglucose isomerase (pgi), which carries a large flux in wild type, *Flux Transformer* inferred zero flux in the mutant from the labeling data alone, matching the published MFA solution. For deleted enzymes of the oxidative pentose phosphate and Entner-Doudoroff pathways, it returned decreased but non-zero values (Fig. 5c).

Since the identity of a genetic perturbation is often known, we next asked whether this information could be used to refine *Flux Transformer* predictions. To this end, we used *FBApro*^**23**^ to project *Flux Transformer* fluxes onto a constraint-based metabolic model in which the deleted reaction was constrained to carry zero flux (*Flux Transformer* + FBApro; see Methods). For deletion of 6-phosphogluconate dehydrogenase (gnd), this projection improved agreement with experimental labeling data from R^2^ = 0.83 to 0.98 (Fig. S4), indicating that perturbation-specific biochemical constraints can provide complementary information beyond the learned prediction alone. In contrast, projection had little effect for deletion of 6-phosphogluconate dehydratase (edd) or Entner-Doudoroff aldolase (eda), for which Flux Transformer already achieved near-perfect agreement with experimental data (R^2^ ≈ 0.98; Fig. S4). For zwf and tpiA, incorporating the perturbation constraint did not improve predictions because the underlying network model failed to capture the relevant metabolic adaptations. Consistent with this, additional network modifications were required to fit these strains in the original study^22^. Thus, Flux Transformer with FBApro can refine predictions or identify gaps in biochemical knowledge.

## Discussion

Quantifying metabolic flux in cells is valuable for a wide range of metabolism-related research^24– 26^. Yet despite decades of method development^26–30^ and publications of the methods and associated tutorials^1,31,32^, metabolic flux analysis remains a specialist technique. This is partly because flux analysis software has traditionally been closed source (e.g., INCA), with open-source engines such as 13CFLUX (v3) only recently available^33^. In addition, MFA tools involve computationally intensive non-linear optimization, do not share common standards and thus make it difficult to share models and workflows, and rely on manual data entry rather than standard mass spectrometry data formats^34–36^. Beyond the software, the analysis itself demands expert judgment at several steps. For example, researchers must decide how to weigh the contributions of the different data types in the cost function.

To lower these hurdles, we developed *Flux Transformer*, a new type of AI model that uses an attention mechanism to weigh the experimental data autonomously and to compute fluxes directly. The model integrates diverse available experimental data and allows researchers to compute flux estimates even when data are sparse. Crucially, *Flux Transformer* models are pre-trained and validated by experts, enabling routine biologist to perform metabolic flux analysis simply by uploading their data to a website (link) which returns accurate fluxes in seconds.

Pre-trained models completely change the economics of the most expert-dependent step. The effort of building and validating a *Flux Transformer* model is spent once, during training, and every subsequent analysis with that model is essentially free. Beyond releasing our pretrained models as a community resource, we provide a site where validated models can be deposited, downloaded and run directly in the browser. Sharing models this way also means that improvements propagate. We anticipate that flux analysis will thus become a cumulative, community effort, as curated genome-scale reconstructions already are.

Because *Flux Transformer* infers flux distributions in seconds and without expert intervention, it makes flux analysis feasible at scales that were previously out of reach: Hundreds of engineered strains, or large panels of conditions. We anticipate that, as with protein structure, the availability of fast and accessible prediction will change not only how fluxes are measured, but how often the question is asked.

### Limitations

*Flux Transformer*, as presented here, infers fluxes at metabolic and isotopic steady state in cultured cells, from bacteria and fungi to plant embryos and mammalian cell lines. Studies where fluxes are inferred from non-stationary labeling data are not yet covered. Because Flux Transformer learns from simulations, extending it to these settings should be feasible by generating suitable training data.

A model is further specific to the organism, the analytical platform (GC- or LC-MS), and the tracers it was trained on, so a new tracer or organism requires retraining. This takes less than an hour on a single GPU and is done once. Since the tracers in common use are few, a small number of models covers most experiments. Quantitative predictions require measured uptake, secretion, and growth rates, since labeling alone constrains only relative fluxes.

Some fluxes are not determined by the labeling data, particularly in larger networks with unidentifiable loops. This is a property of the experiment and the network, not of the method, and applies equally to traditional MFA. *Flux Transformer* reports such fluxes with wide confidence intervals, so gaps are visible immediately. Its speed then makes it attractive to combine *Flux Transformer* with simulation and analysis modalities that bring in orthogonal knowledge, such as, the integrating of transcriptomic^37^ and proteomic data^38^, or thermodynamic^39^ and resource allocation^40^ constraints.

The *Flux Transformer* models presented here are restricted to cultured cells or plant embryos and do not yet cover organismal metabolism. The efficiency gains observed here provide optimism that related methods will eventually enable flux analysis at organismal scales.

## Data and code availability

The source code, datasets and network models for *Flux Transformer* are available at https://github.com/weilandtd/flux-transfomer under the APACHE-2 license. The four pretrained and validated models (*E. coli, S. cerevisiae, I. orientalis*, pennycress embryos, and mouse T cells) can be downloaded from GitHub.

## AI disclosure

We used Claude (Anthropic) to assist with writing the manuscript and the software for this study. We reviewed and verified every output, and we take full responsibility for the content of the manuscript and the software.

## Acknowledgements

We thank members of the Rabinowitz and Singh labs for discussions. We specifically want to thank Ariel Brunner for helpful comments and valuable insights on flux analysis. We also thank Ron Milo valuable feedback on figures. This work was funded by Department of Energy Center for Advanced Bioenergy and Bioproducts Innovation (U.S. Department of Energy, Office of Science, Biological and Environmental Research Program under Award Number DE-SC0018420) supporting J.D.R., D.R.W., and J.S. This work is further supported in part by the National Institute of General Medical Sciences of the National Institutes of Health under grant number R35GM158278 (to M.S.), and by the Princeton Laboratory for Artificial Intelligence. Research in the Rabinowitz laboratory was supported by NIH Grant R01CA163591 and Ludwig Cancer Research. Additional support was provided by the University of Pennsylvania Diabetes Research Center-Princeton Inter-Institutional Metabolomics Core (NIH Grant P30DK19525) and the Rutgers Cancer Institute of New Jersey Cancer Center Support Grant (NIH Grant P30CA072720). Any opinions, findings, conclusions or recommendations expressed in this publication are those of the author(s) and do not necessarily reflect the views of the US DOE or NIH.

## Author contributions

D.R.W. and J.D.R. conceived the study. D.R.W. and M.T.T. developed the model architecture, designed and performed the simulation studies, generated the training and benchmarking datasets, and applied the framework across diverse metabolic network models. D.R.W. and M.T.T. led the implementation of the methods and models (software). R.O. designed and led the implementation of the web portal. J.S. contributed plant metabolic models and domain expertise that informed model development and evaluation. D.R.W., J.D.R., and M.S. supervised the research. D.R.W., M.T.T., J.D.R., and M.S. wrote the manuscript with input from all authors.

## Declaration of interests

The authors declared no competing interests.

## Methods

### Isotope tracing datasets

Five isotope-tracing datasets were curated from the literature, including measurements obtained using liquid chromatography-mass spectrometry (LC-MS) and gas chromatography-mass spectrometry (GC-MS). Metabolite isotopologue distributions were measured by LC-MS in wild-type *Escherichia coli*, yeasts (*Saccharomyces cerevisiae, Issatchenkia orientalis*), and activated and naïve T cells across multiple experimental conditions and by GC-MS in pennycress (*Thlaspi arvense L*) embryos. A separate GC-MS dataset comprising wild-type and mutant *E. coli* strains was included for comparative analysis, with the mutant strains carrying genetic perturbations affecting central carbon metabolic pathways.

### Isotope labeling models

In total, four isotopic tracing models were collected from literature^16,18,19^ and translated into the common COBRA json format, using the *annotation* field to specify atom mapping between reactants. To account for compartmental mixing introduced during metabolite extraction, the yeast and mouse models were modified to include reactions that pool cytosolic and mitochondrial metabolites^41,42^. To account changes in cell composition in yeast and mouse cells, the biomass reaction was reformulated such that production rates of the individual macromolecule classes (DNA, RNA, lipids, and protein) are independent.

To recapitulate the isotopologue distributions observed in GC-MS measurements, models were modified to explicitly represent metabolite fragmentation occurring during ionization and detection. Fragment-specific atom mappings were incorporated into the measurement model, based on a literature generated library for metabolite fragments (Data S2), enabling direct simulation of the isotopologue distributions associated with experimentally quantified fragment ions.

### Labeling simulations

Stable-isotope labeling was simulated from steady-state balanced flux distributions using the elementary metabolic unit (EMU) algorithm^5^. These flux distributions were generated by an artificially centered hit and run (ARCHR) algorithm^43^, that randomly sampled steady-state balanced fluxes within physiological bounds using a flux balance model of the reaction network^44^. To compute labeling efficiently in large reaction networks, a GPU compatible version of EMU was implemented using PyTorch^45^. This implementation further directly reads COBRA json models, annotated with atom transitions. All codes for this implementation are available in the GitHub repository (https://github.com/weilandtd/flux-transfomer).

### Measurement tokenization

*Flux transformer* represents each experimental measurement as an individual token that is processed by the attention layers. The input consists of two measurement types: isotopologue measurements and reaction rate measurements. The identity of an isotopologue measurement is defined by three components: the metabolite, the mass shift (M+i), and the tracer used in the experiment. In contrast, the identity of a reaction rate measurement is defined by its corresponding reaction. For metabolites and reactions, these identities are defined from the reaction network used to generate the training data, independent of which measurements are available in a particular experiment.

Isotope-labeling measurements were represented by three categorical attributes: the tracer, measured metabolite, and isotopologue mass shift. All tracers, metabolites, and mass shifts represented in the dataset were assigned unique integer identifiers beginning at zero within their respective categories. Because these identifiers serve only as categorical labels and do not imply an intrinsic ordering, they were transformed into bounded radial embeddings rather than supplied directly to the model. For an identity with label *k* among *N* possible identities of a given type, the embedding was defined as:

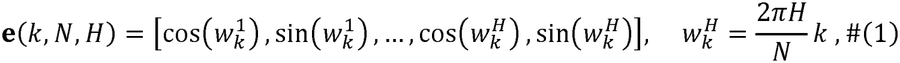

where *H* is the number of embedding frequencies. Each measured isotopologue fraction *x* is represented as a four-dimensional vector

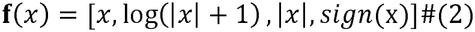

The representation of an isotope-labeling measurement was then constructed by concatenating the embeddings of its constituent attributes and it associated value,

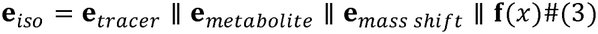

where ∥ denotes vector concatenation, and 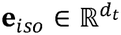. The resulting vector dimensionality *d*_t_ is a function of the embedding frequencies *H* chosen for the tracer, metabolite and mass shift embeddings and may be tuned to represent larger metabolic models with more metabolites.

Each reaction in the network was assigned a unique integer identifier beginning at zero and encoded using the same bounded radial embedding formulation as the isotopologue fraction. For a measured reaction rate *v*, we construct the resulting reaction embedding as,

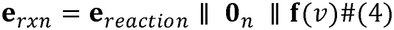

where *f*(*v*) encodes the measured rate using Equation (2), and is made to be of the same dimensionality *d*_*t*_ as the isotope labeling embeddings, by padding the vector with *n* zeros, i.e., 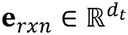. The resulting tokens are provided jointly to the attention layers.

### Choosing free fluxes

To efficiently learn how measurement tokens relate to the flux distribution, *Flux transformer* reduces the dimensionality of the output by considering a metabolic steady state. At metabolic steady state, intracellular metabolite concentrations remain constant, and the reaction fluxes therefore satisfy the mass-balance constraint

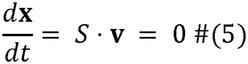

where *S* ∈ ℝ^*m*×*n*^ is the stoichiometric matrix and **v** ∈ ℝ*n* is the vector of reaction fluxes. The set of fluxes satisfying this constraint lies in the null space of *S*. If *S* has rank *r*, this null space has dimension *d* = *n* ™ *r*. Let the matrix *Z* ∈ ℝ*n*×*d* have columns that form a basis for this null space. Every mass balanced flux distribution **v** can then be written as

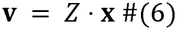

for some **x** ∈ ℝ*d*. Flux Transformer predicts the *d* coordinates of 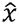 and reconstructs the full distribution as 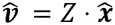. Then since

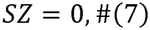

flux predictions 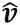 obey metabolic state steady for any 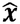.

The null-space basis is not unique. A standard basis obtained from an QR decomposition generally represents each reaction as a combination of several null-space coordinates. Instead *Z* was constructed so that each measured reactions *t* ∈ *T* is predicted directly as a free flux. Let *j*_*t*_ denote the position of reaction *t* in the vector of free fluxes. Then

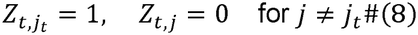

To construct such a basis, *r* independent rows are identified from the stoichiometric matrix by QR decomposition producing a row reduced stoichiometric matrix *S*_*r*_ ∈ ℝ^*r*×*n*^. Then pivoted QR decomposition on the unmeasurable columns *l* ∉ *T*, identifies a robust dependent set *D*, of fluxes that excludes the measurable reactions. The set of free fluxes was then defined by exclusion, i.e. *F* are all *f* ∉ *D*, and encompassed measurable rates by construction.

This allows to partition *S*_*r*_ into free and dependent columns, *S*_*D*_ ∈ ℝ^*r*×*r*^ and *SF* ∈ ℝ^*r*×*d*^ respectively. The steady-state constraint can then be written as:

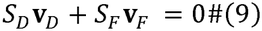

and can be solve for the dependent fluxes as

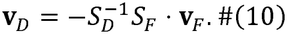

Let 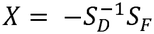, then the constrained basis *Z* can then be directly constructed as

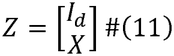

where *I*_d_ is the identify of size *d* = *n* − *r*, and *X* ∈ ℝ^*r*×*r*^ is computed from *S*_*D*_ and *S*_*F*_. The resulting basis therefore provides a *d*-dimensional representation of steady-state fluxes in which reaction rates that can be measured are explicitly represented whenever possible.

### AI-model architecture

The Flux Transformer maps a variable number of experimental measurements to a fixed set of free flux coordinates. Each measurement is first represented as a token embedding encoding the associated experimental information.

Tokenized measurements were processed jointly using a stack of transformer self-attention layers^13^. Given *N* measurement tokens,

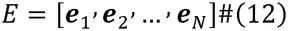

the model computes query, key, and value representations from the learned projection matrices *WQ*, 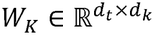 and 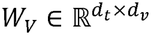:

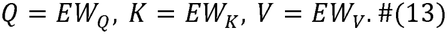

where *Q*, 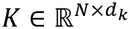 and 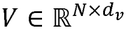. Self-attention is then computed as

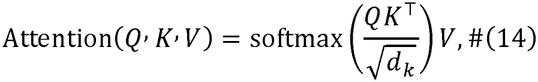

Here, *d*_*k*_ and *d*_*v*_ denote the dimensionality of the query/key and value representations, respectively. The factor 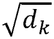 scales the dot-product similarity scores to maintain a stable magnitude as the embedding dimension increases. The resulting attention weights determine the contribution of each measurement token to the updated representation of each token, allowing information from distinct experimental measurements to be integrated before flux prediction. Each layer applies *h* such attention head in parallel, each with its own projection matrices and *d*_*k*_ = *d*_*v*_ = *d*_*t*_/*h*, and the head outputs are concatenated and projected back into dimensionality *d*_*t*_,

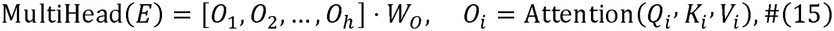

with the learned output projection 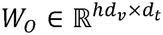. The resulting contextualized measurement representations,

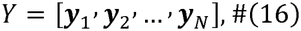

integrate information across measurements originating from different tracers, metabolites, and experimental conditions.

To predict metabolic fluxes, the model employs a cross-attention decoder comprising *L* layers that maps the contextualized measurement representations to the *d* free coordinates of the flux null space. A set of learned query embeddings,

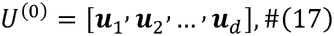

is initialized, with one query associated with each free flux coordinate. During cross-attention, these query embeddings attend to the complete set of measurement representations,

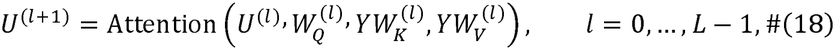

allowing each query to assign its own weights to the measurements when predicting its corresponding free reaction rate. Because all queries attend to the same measurement set, different flux coordinates can selectively utilize distinct combinations of isotopic and reaction rate measurements.

The final query representations are independently projected to a scalar value for each free flux coordinate:

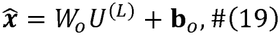

where the output biases **b**_*o*_ ∈ ℝ^*d*^ and weights *W*_*o*_ ∈ ℝ^*d*×*t*^ are learned during training. These coordinates are subsequently transformed into the full flux vector through the null-space parameterization described above.

### Loss function and training

The Flux Transformer was trained on simulated data to predict the full flux state from isotopologue distributions and measurable reaction rate. Errors were computed on the free fluxes *y* which were computed from the fluxes *v* using the basis representation *Z*:

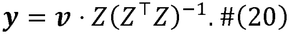

We found that a loss function based on the variance-normalized mean square error performed better than standard mean squared error:

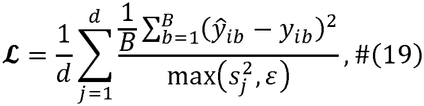

where *B* is the minibatch size, *d* is the number of refluxes, 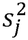 is the training-set variance for each free flux, and *ε* = 0.1 is a lower bound on that variance.

For each organism, 10,000 simulated samples were split 90:10 into training and validation sets. Models were optimized with Adam (learning rate 10^-3^, batch size 256, gradient clipping at 1.0), with a 5-epoch linear warmup and learning-rate halving when the validation loss plateaued. Training stopped early once the validation loss had not improved for 10 epochs, and the best validation checkpoint was kept. To make the model robust to experimental error, each minibatch was augmented with Gaussian noise on the isotopologue distributions (0.01 absolute) and on the measured fluxes (5% relative). Each input token was also masked with probability 0.2. An ensemble of 20 independently initialized models was trained per organism.

### Uncertainty estimates

Inference uncertainty has two sources: noise in the measurements, and disagreement between the ensemble members. We captured both by bootstrapping the measurements through the full ensemble. In each bootstrap draw (B = 200), we picked one biological replicate per tracer at random, and sampled each measured exchange flux within its reported error, without changing its sign. Every ensemble member predicted fluxes for every draw. The reported value is the ensemble prediction at the mean of the replicates. The 95% interval is the percentile interval over all member predictions from all draws reflecting measurement and model uncertainty.

### Flux transformer with FBApro

We integrated knock-out information into *Flux Transformer* solutions by applying *FBApro*^23^. This method maps a predicted flux vector 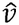 to the closest steady-state flux vector by an explicit orthogonal projection onto the null space of the stoichiometric matrix *S*, while holding a set of chosen reactions *K* at fixed values. Here, *K* is the set knocked-out reactions fixed at zero flux.

Enforcing a knockout forces the network to reroute flux, and the projection determines which reactions absorb this change. In its original formulation, *FBApro* weights every reaction equally, so the change is spread across all reactions, including those the *Flux Transformer* predicts confidently from the measurements. We instead used a weighted projection, so that fluxes with small uncertainty are constrained more than fluxes with large uncertainty. This weighted projection can be simply constructed by weighting the input and output of the *FBAproFixed* operator using Λ the diagonal matrix of all standard deviations *σ*_*i*_:

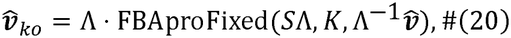

where *K* is the set of knocked-out reactions, and 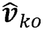 is the projection of the predicted flux vector 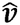 with the knocked-out fluxes set to zero. The resulting 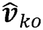 solves the weighted least-square problem of finding the steady state flux vector closest to 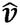 with each squared deviation weighted by 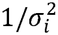, and the knocked-out fluxes fixed at zero.

### Traditional metabolic flux analysis

Benchmark metabolic flux analysis (MFA) was performed using 13CFLUX v3.0 and INCA v2.5. For each isotope-tracing dataset, fluxes were estimated using the corresponding published network reconstruction and measurement model. Optimization and confidence interval estimation were performed using the default workflows implemented in each software package unless otherwise specified.

### Evaluation metrics

To evaluate how well the different flux identification methods recall the true (input) fluxes, given a model and a set of experimental data, for each reaction *j*, we computed the coefficient of determination 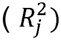 between the ground-truth observed *v*_*ij*_ and predicted flux 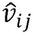 across *i* validation (N=1000) samples.

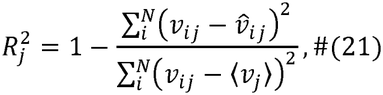

were ⟨ *v*_*j*_ ⟩ is the mean of the observed flux for reaction . Similarly, we computed condition specific coefficients of determination (R^2^) for *Flux Transformer* inferred fluxes against the expert curated MFA fluxes from literature

We also assessed overall performance across species by computing the mean normalized error on the validation dataset as

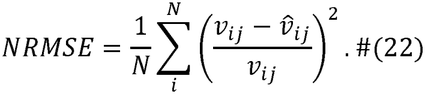

Runtime of the different algorithms was assessed as the mean time each algorithm requires to compute a single flux distribution from a single set of input data.

## Notes

### Competing Interest Statement

The authors have declared no competing interest.

