## Supplemental Figures for "*Flux Transformer* learns intracellular metabolic fluxes from isotope labeling and measured reaction rates"

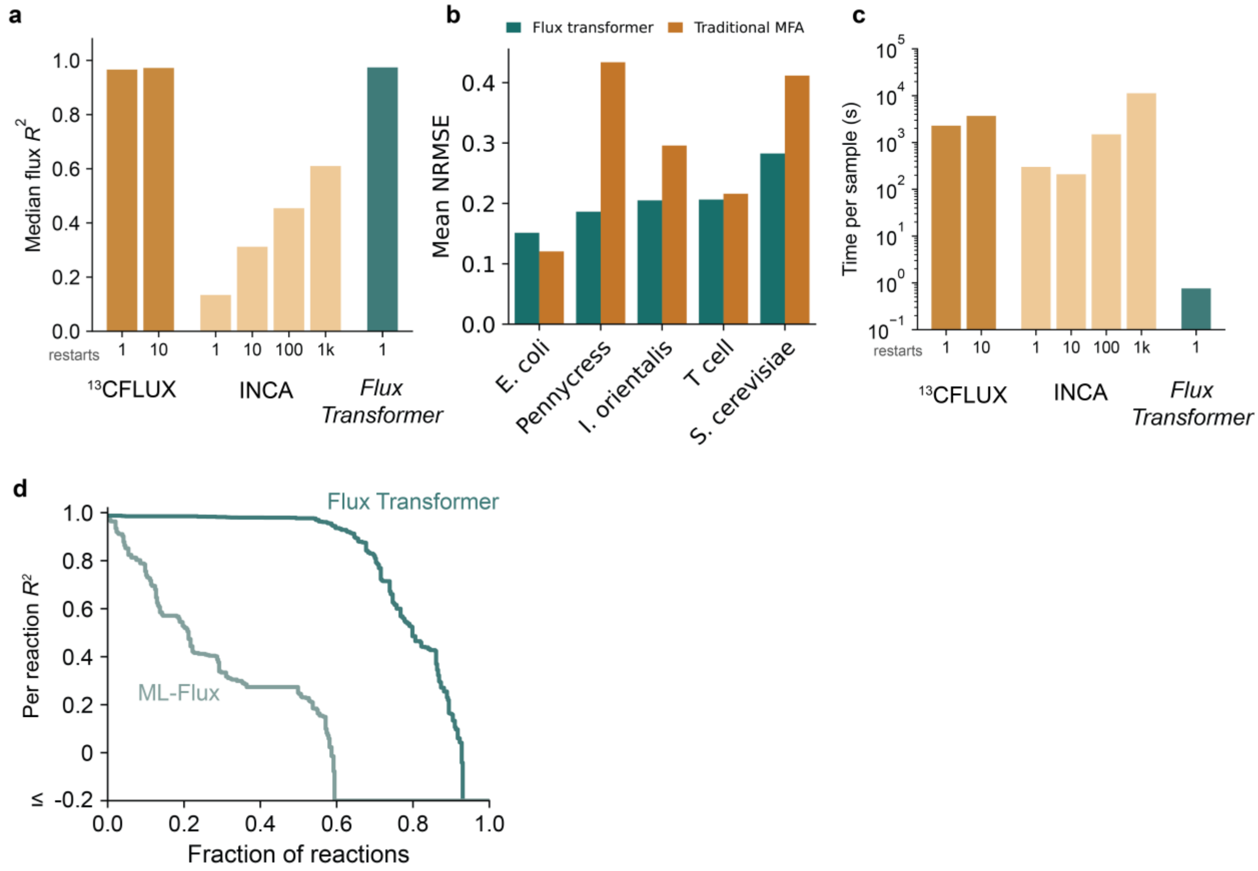


**Figure S1: *Flux Transformer* achieves accurate flux predictions with substantially reduced computational cost.**

(a) Median reaction-level prediction accuracy ($R^2$) in *I orientalis* achieved by traditional metabolic flux analysis methods ($^{13}$CFLUX and INCA) as a function of the number of optimization restarts, compared with *Flux Transformer*.

(b) Mean normalized root-mean-square error (NRMSE) of flux inferred on noisy data compared to "ground truth" across five different organisms for *Flux Transformer* and traditional metabolic flux analysis.

(c) Average computation time per sample for traditional metabolic flux analysis methods ($^{13}$CFLUX and INCA) with varying numbers of optimization restarts, compared with *Flux Transformer*. Time is shown on a logarithmic scale.

(d) Distribution of per-reaction flux agreement ($R^2$) for Flux Transformer and ML-Flux. Reactions are ranked by prediction accuracy. ML-Flux exhibits substantially lower and more variable performance.

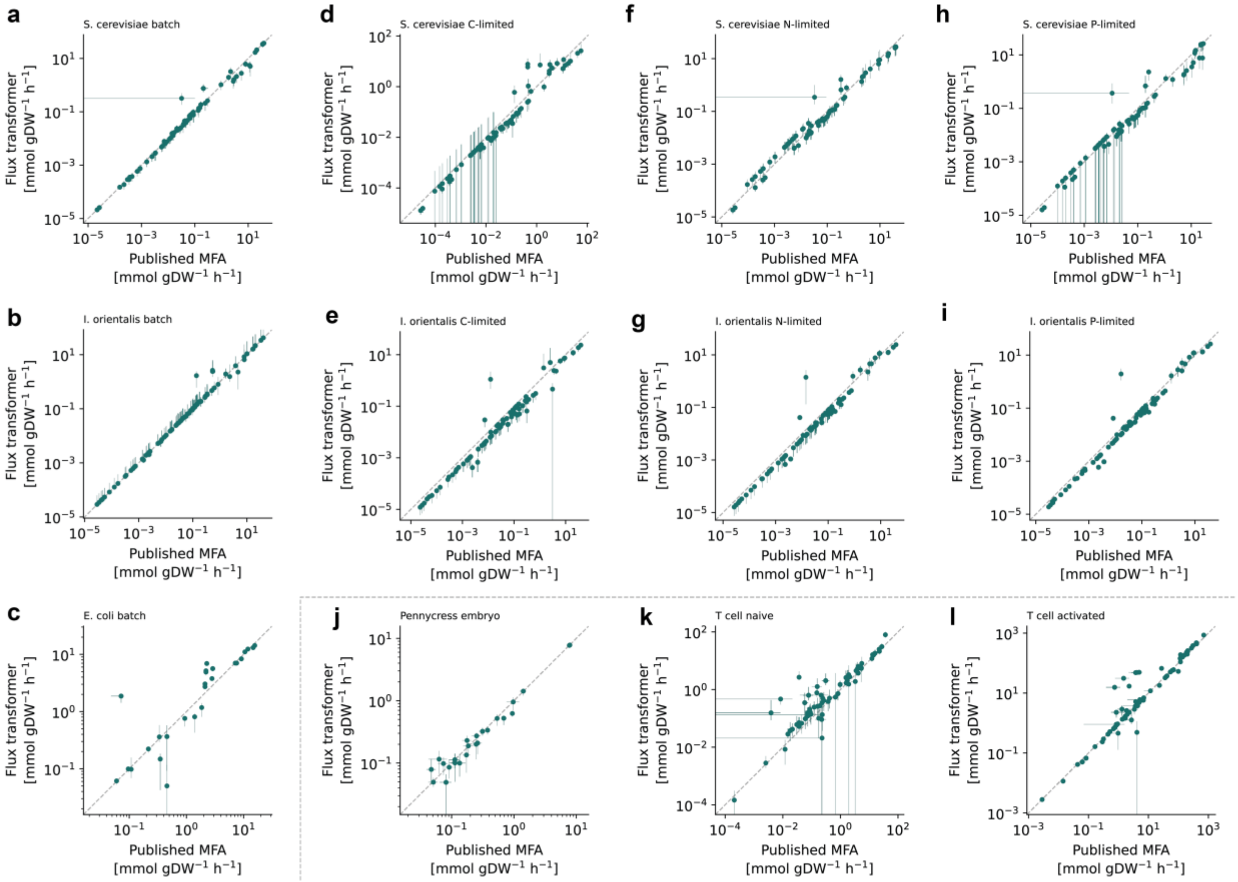


**Figure S2: Flux Transformers accurately reproduce published metabolic fluxes across diverse organisms, and growth conditions.**

(a–l), Flux Transformer predictions versus published metabolic flux analysis (MFA) flux estimates for *S. cerevisiae* batch (a), *I. orientalis* batch (b), *E. coli* batch (c), *S. cerevisiae* carbon-limited (d), *I. orientalis* carbon-limited (e), *S. cerevisiae* nitrogen-limited (f), *I. orientalis* nitrogen-limited (g), *S. cerevisiae* phosphorus-limited (h), *I. orientalis* phosphorus-limited (i), pennycress embryos (j), naive T cells (k), and activated T cells (l). Each point represents a reaction flux, error bars indicate uncertainty estimates from MFA, and dashed lines indicate the identity relationship. Flux values are shown on logarithmic scales.

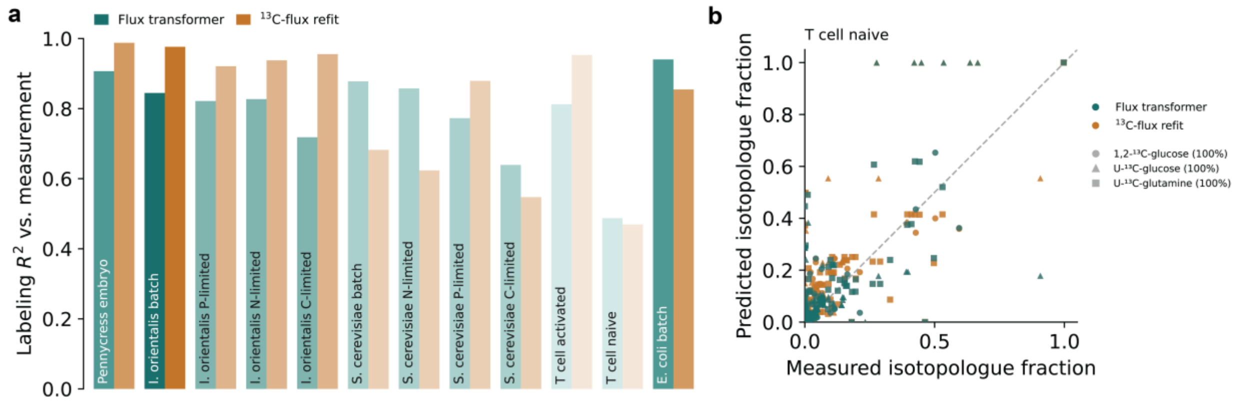


**Figure S3: Flux Transformer prediction accuracy mirrors the fit achievable by expert-curated metabolic flux analysis.**

(a) Agreement between predicted and measured isotopic labeling patterns for *Flux Transformers* and MFA refits that were constrained closely to expert curated fluxes. Bars report $R^2$ across all measured isotopologue fractions within each dataset. Datasets showing lower *Flux Transformer* performance also exhibited lower agreement for MFA refits, suggesting that prediction accuracy is limited by the information content of the labeling data and the ability of the underlying model to explain the measurements.

(b) Predicted versus measured isotopologue fractions for naive T cells. Points represent individual isotopologue measurements from [1,2-$^{13}$C]glucose, [U-$^{13}$C]glucose, and [U-$^{13}$C]glutamine tracer experiments. The dashed line indicates the identity relationship

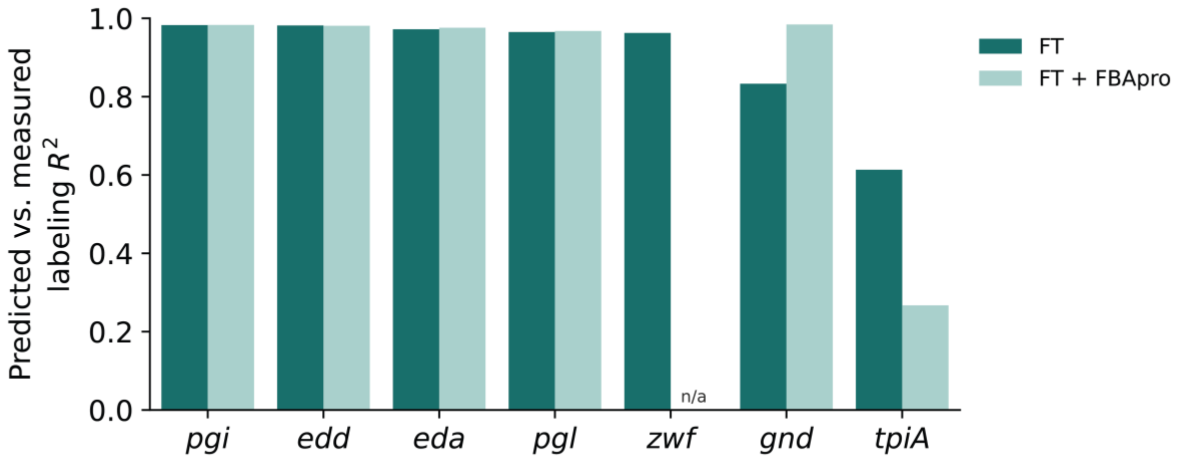


**Figure S4: Constraint-based refinement can improve labeling predictions when perturbation-induced metabolic rewiring is captured by the biochemical model.**

Comparison of isotopologue prediction accuracy for Flux Transformers (FT) and Flux Transformers combined with FBApro (FT + FBApro) across *E. coli* reaction knockdown strains. FT + FBApro denotes Flux Transformer predictions projected onto a constraint-based metabolic model in which the deleted reaction is constrained to zero flux. Bars report agreement between predicted and measured isotopic labeling patterns ($R^2$). Incorporating perturbation information improved prediction accuracy for the *gnd* knockdown, suggesting FBApro can provide complementary information when the metabolic consequences of a perturbation are accurately represented. In contrast, projection did not improve predictions for *zwf* and *tpiA*, because biochemical adaptations required to explain these perturbations are absent from the network model.
